# Genome sequencing and assessment of plant growth-promoting properties of a *Serratia marcescens* strain isolated from vermicompost

**DOI:** 10.1101/288084

**Authors:** Filipe P. Matteoli, Hemanoel Passarelli-Araujo, Régis Josué A. Reis, Letícia O. da Rocha, Emanuel M. de Souza, L. Aravind, Fabio L. Olivares, Thiago M. Venancio

## Abstract

Plant-bacteria associations have been extensively studied for their potential in increasing crop productivity in a sustainable manner. *Serratia marcescens* is a Gram-negative species found in a wide range of environments, including soil. Here we describe the genome sequencing and assessment of plant-growth promoting abilities of *S. marcescens* UENF-22GI (SMU), a strain isolated from mature cattle manure vermicompost. *In vitro*, SMU is able to solubilize P and Zn, to produce indole compounds (likely IAA), to colonize hyphae and counter the growth of two phytopathogenic fungi. Inoculation of maize with SMU remarkably increased seedling growth and biomass under greenhouse conditions. The SMU genome has 5 Mb, assembled in 17 scaffolds comprising 4,662 genes (4,528 are protein-coding). No plasmids were identified. SMU is phylogenetically placed within a clade comprised almost exclusively of environmental strains. We were able to find the genes and operons that are likely responsible for all the interesting plant-growth promoting features that were experimentally described. Genes involved other interesting properties that were not experimentally tested (e.g. tolerance against metal contamination) were also identified. The SMU genome harbors a horizontally-transferred genomic island involved in antibiotic production, antibiotic resistance, and anti-phage defense via a novel ADP-ribosyltransferase-like protein and possible modification of DNA by a deazapurine base, which likely contributes to the SMU competitiveness against other bacteria. Collectively, our results suggest that *S. marcescens* UENF-22GI is a strong candidate to be used in the enrichment of substrates for plant growth promotion or as part of bioinoculants for Agriculture.

## INTRODUCTION

The current projections of the world population growth creates an increasing pressure for the adoption of intensive farming, often resulting in reduced soil fertility, eutrophication of aquatic and terrestrial environments and destruction of the biodiversity (Bhardwaj et al. 2014). Moreover, conventional agriculture settings often generate large volumes of organic wastes, constituting a major source of environmental pollution due to rejection or incineration (Angulo et al. 2012). Sustainable strategies to minimize these problems have been investigated worldwide, particularly in developing countries like Brazil, where agriculture plays a major role in the balance of trade (Olivares et al. 2017).

Composting and vermicomposting are widely known techniques that involve the stabilization of organic materials, with a concomitant sustainable production of valuable soil amendments (Quagliotto et al. 2006). Classical composting is defined as the biological decomposition of organic wastes in an aerobic environment carried out by microorganisms (Sim and Wu 2010), while vermicomposting also involves earthworms that promote aeration and help in waste stabilization by fragmenting the organic matter and boosting microbial activity (Domínguez et al. 2003). Vermicomposting has a thermophilic stage, promoted by a thermophilic bacterial community that drives the most intensive decomposition. This stage is followed by a mesophilic maturation phase that is largely mediated by earthworms and associated microbes (Pathma and Sakthivel 2012). Vermicomposted material holds greater amounts of total phosphorus (P), micronutrients and humic acid substances than the original organic material. In general, vermicomposts are considered a safe, cheap and rich source of beneficial microorganisms and nutrients for plants (Hashemimajd et al. 2004). Further, bacteria isolated from vermicompost typically display greater saprophytic competence than those intimately associated with plants. From a biotechnological perspective, the microbial survival and activity in the absence of a host plant represent an ecological advantage that can be used as a strategy to enrich substrates with nutrients, boosting plant growth and development (i.e. plant substrate biofortification) (Busato et al. 2012; Busato et al. 2017).

Plants often benefit from mutualistic interactions with plant growth-promoting rhizobacteria (PGPR) (Ma et al. 2016). PGPR can promote plant growth by various mechanisms, such as: 1) mitigation of abiotic stresses such as metal phytotoxicity (Glick 2010), water or salinity stress (Rho et al. 2017); 2) activation of defense mechanisms against phytopathogens (Hol et al. 2013); 3) directly attacking pathogens (Liu et al. 2017); 4) biological nitrogen fixation (da Costa et al. 2014); 5) solubilization of mineral nutrients (e.g. P and zinc, Zn) (Oteino et al. 2015); 6) production phytohormones (Ortíz-Castro et al. 2009) and; 7) secretion of specific enzymes (e.g., 1-aminocyclopropane-1-carboxylate deaminase) (Sarkar et al. 2017). Due to their interesting beneficial effects, there is a growing market for PGPR biofertilizers (Balasubramanian and Karthickumar 2017), which are based on bacteria of various genera, such as *Azospirillum*, *Bacillus* and *Azotobacter* (Bashan et al. 2014). A notable example of successful application of PGPR in agriculture is the soybean (*Glycine max* L.) production in Brazil, in which the development and use of an optimized consortium of different strains of *Bradyrhizobium* sp. (Souza et al. 2015) led to very high productivity levels at significantly lower costs due to the virtually complete replacement of nitrogen fertilizers (Chang et al. 2015).

Knowledge of PGPR genomic content and plant interaction mechanisms has increased with the progress of second-generation sequencing technologies (MacLean et al. 2009), which also allowed a number of comparative genomics studies. In a large-scale comparative analyses of alpha, beta and gamma-proteobacteria, Bruto et al. found no set of plant beneficial genes common to all PGPR, although the presence of certain genes could reflect bacterial ecological type, such as the presence of *ppdC* (involved in auxin biosynthesis) exclusively in endophytic strains of *Azospirillum* and *Bradyrhizobium* (Bruto et al. 2014). *Bacillus amyloliquefaciens* subsp. plantarum FZB42 is a clear example of how genome mining strategies can uncover the genetic basis of plant-growth promoting capacity of a PGPR (Paterson et al. 2016). After promising results on auxin (Idris et al. 2007) and phytase (Makarewicz et al. 2006) production *in vitro*, genome analysis also uncovered the molecular basis of how this strain exerts its antifungal (Koumoutsi et al. 2004), antibacterial (Wu et al. 2015) and nematicidal activities (Liu et al. 2013). Another important example of genomic analysis of a PGPR is that of *Herbaspirillum seropedicae* SmR1, in which genes associated with nitrogen fixation and plant colonization were elegantly investigated (Pedrosa et al. 2011).

*Serratia marcescens* is a Gram-negative and rod-shaped bacteria that has been proposed as a PGPR due to its P solubilization properties (Ben Farhat et al. 2009; Tripura et al. 2007), chitinase activity (Vaikuntapu et al. 2016) and prodigiosin-mediated insect biocontrol (Suryawanshi et al. 2015). *S. marcescens* has been described in association with several plants, such as cotton (*Gossypium hirsutum*) and maize (*Zea mays*) (McInroy and Kloepper 1995), rice (*Oryza sativa*) (Gyaneshwar et al. 2001) and pinus (*Pinus pinaster*) (Vicente et al. 2016). *S. marcescens* FS14 (isolated from *Atractylodes macrocephala*) was shown to exert antagonistic effects against phytopathogenic fungi and genomic sequencing revealed the presence of an interesting pattern of secretion systems (Li et al. 2015). Some *S. marcescens* isolates have also been reported as opportunistic pathogens (Mahlen 2011) and most comparative genomics studies of this species focused exclusively on its clinical relevance (Moradigaravand et al. 2016). A comparative analysis of insect and clinical *S. marcescens* isolates revealed a substantial genetic diversity, as supported by a relatively low intra-species average nucleotide identity (ANI) of 95.1%. Further, a type II secretion system, often related to virulence (Korotkov et al. 2012), was found in the clinical but not in the insect strain (Iguchi et al. 2014).

Here we report a comprehensive characterization of *S. marcescens* UENF-22GI (SMU), a strain that has been shown to be abundant in mature cattle manure vermicompost, from where it was isolated. We performed a series of *in vitro* and *in vivo* experiments that show SMU’s ability to solubilize P and Zn, to synthesize indole compounds (likely the auxin indole acetic acid, IAA) and to counter the growth of phytopathogenic fungi. Inoculation with SMU substantially increased maize growth and biomass under greenhouse conditions. Given its promising results as a PGPR, we sequenced its genome and carefully identified the genetic basis of these and other important features. The SMU genome also harbors an interesting horizontally-transferred genomic island involved in production of a peptide antibiotic and phage resistance via modification of DNA by a deazapurine base, which probably contributes to its competitiveness against other microorganisms. Further, phylogenetic reconstructions placed SMU in a clade that mainly comprises non-clinical *S. marcescens* isolates. Collectively, our results strongly indicate that SMU is a good candidate to be used in inoculant formulations or as part of a strategy for biological enrichment of plant substrates.

## RESULTS AND DISCUSSION

### Identification of the isolate

During the initial characterization of abundant culturable bacteria from the mature cattle vermicompost, we identified a notorious pigmented bacterium that was preliminarily characterized as a *S. marcescens* by colony morphology, microscopy and 16S rRNA sequencing. This isolate was named *Serratia marcescens* UENF-22GI (SMU). SMU was tested for a series of plant growth-promotion traits and, given the promising results, submitted it to whole-genome sequencing and comparative analysis.

### *In vitro* solubilization of P and Zn and synthesis of indole compounds by SMU

We explored the capacity of SMU to solubilize P and Zn *in vitro*, as the availability of these elements is often a limiting factor in crop production (Mehra et al. 2017). We used the formation of a halo as a positive result for the solubilization of P and Zn, which are essential nutrients for bacterial growth. The halo and colony dimensions were also used to calculate a solubilization index (SI), which is a useful metric to estimate the P and Zn solubilization capacities. Our results clearly show that SMU solubilizes P and Zn *in vitro*, with SI values of 2.47 ± 0.22 and 2.11 ± 0.47, respectively (Figure 1a and b). We have also used a quantitative approach to measure P solubilization using two distinct inorganic P sources: calcium phosphate (P-Ca) and fluorapatite rock P (P-rock). Remarkably, we found that SMU increases the amount of soluble P by 12- and 13-fold with P-Ca and P-rock, respectively (Figure 1c). Because acidification is a common P solubilization mechanism, we have also monitored pH variation and found a striking media acidification using glucose as carbon source, from 7.0 to 3.78 and 3.54 for P-Ca and P-rock, respectively (Figure 1d). Importantly, most P-solubilization screenings are conducted only using Ca-P. However, in most tropical soils, P is typically associated with Fe and Al. The ability of SMU to solubilize P from P-rock is important, as this P source is recommended for organic agricultural systems. Hence, we propose that P-rock and SMU could be used in combination as a P-fertilization strategy for tropical soils.

**Figure 1:**
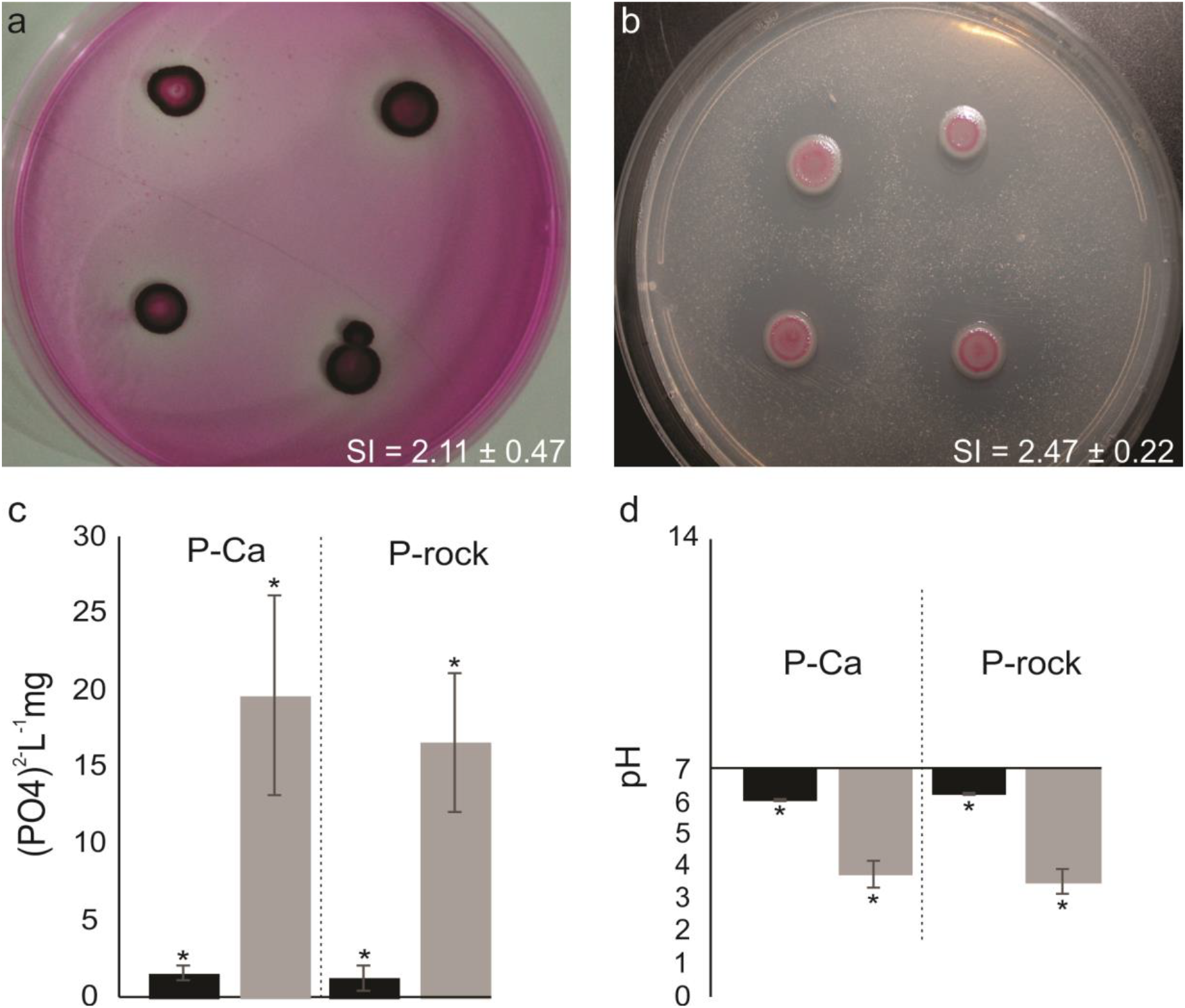
Phosphorus (a) and zinc (b) solubilization assays. Qualitative P and Zn solubilization assays were carried out with Ca_3_(PO_4_)_2_ (P-Ca) and ZnO as substrates, respectively. Halo formation around growing colonies was considered a positive result for solubilization. These results were used to compute the solubilization index (SI), which is the halo diameter divided by the colony diameter. Quantitative P solubilization assays were also performed using P-Ca or fluorapatite rock phosphate (P-rock) in the absence (black bars) or presence (gray bars) of SMU (c). pH variation in the culture media in the absence (black bars) or presence (gray bars) of SMU, indicating that P solubilization is probably driven by acidification (d).

Bacterial production of phytohormones (e.g. indole-3-acetic acid, IAA, an auxin) is considered a major factor in enhancing plant growth (Santoyo et al. 2016). IAA is a primary regulator of plant growth and development. At least four tryptophan-dependent IAA biosynthesis pathways have been identified in Bacteria: indole-3-acetamide (IAM), indole-3-acetonitrile (IAN), indole-3-pyruvic acid (IPyA) and tryptamine (TAM) pathways (Duca et al. 2014; Spaepen and Vanderleyden 2011). Since the IAA biosynthesis genes are also involved in the Ehrlich pathway (degradation of amino acids via transamination, decarboxylation and dehydrogenation), gene presence alone is not sufficient to determine IAA production. Therefore, we tested the ability of SMU to synthesize indole compounds *in vitro* and verified that it is indeed able to produce IAA either in the presence or in the absence of Trp. However, greater IAA levels were observed in the former condition (Figure 2).

**Figure 2:**
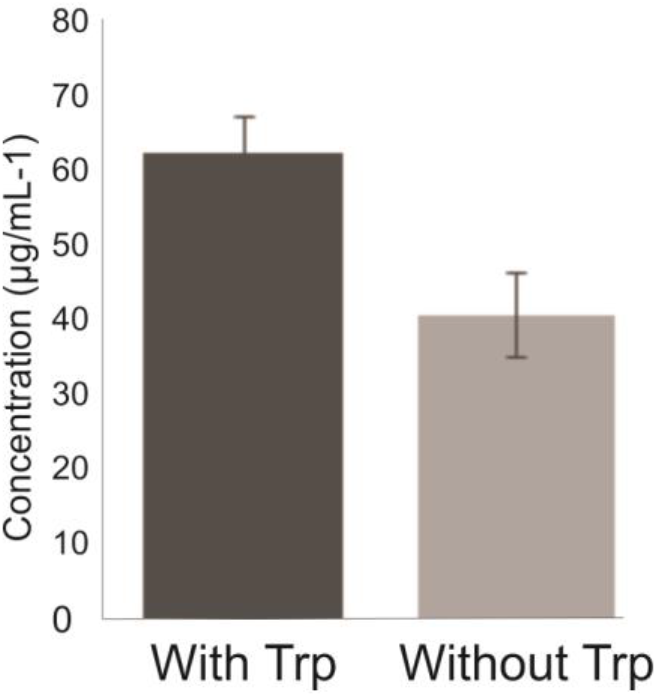
Biosynthesis of indole compounds in the presence and absence of Trp. An aliquot of the SMU inoculum was transferred to Dygs medium with or without tryptophan (100 mg⋅L^−1^) and incubated for 72 h in the dark, at 30 °C and 150 rpm. To evaluate indole synthesis, 150 µL of grown bacteria were transferred to microplates and 100 µL of Salkowski reagent (see methods for details) were added. The plate was incubated for 30 min in the dark and samples analyzed at 492 nm on a spectrophotometer.

### Biofilm formation and biocontrol of phytopathogenic fungi

Many fungi colonize the rhizosphere and must cope with strong competition from soil bacteria. Several of these fungi are phytopathogenic and pose serious risks to agriculture (Schwessinger et al. 2015). In order to assess the antifungal properties of SMU, we performed a dual growth assay and found that SMU counters the growth of *F. oxysporum* and *F. solani* (Figure 3). The strategy deployed by SMU to hinder fungal growth probably involves massive biofilm formation on *Fusarium* hyphae (Figure 3, Figure S1), which probably facilitates the colonization and degradation of fungal cell walls. In addition, there is a conspicuous delineation of the space occupied by *F. solani* by prodigiosin (Figure S1), supporting the previously proposed antifungal activity of this this secondary metabolite (Duzhak et al. 2012).

**Figure 3:**
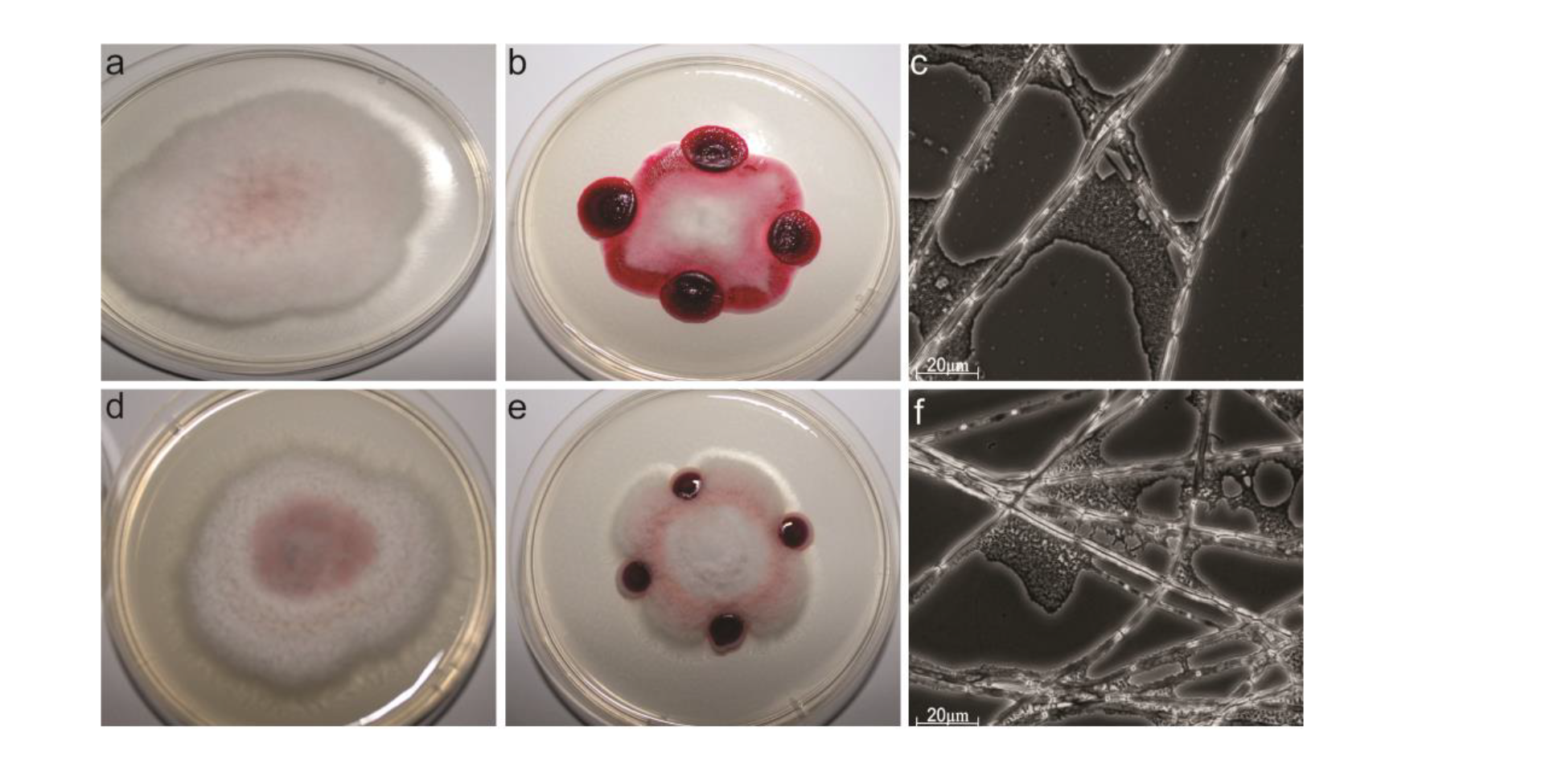
Dual growth assays of SMU and two phytopathogenic *Fusarium* species. Controls were conducted with *F. oxysporum* and *F. solani* grown without SMU (a and d, respectively). In the dual growth assays, SMU was placed in four equidistant regions to the *F. oxysporum* and *F. solani* (b and e, respectively). The adherence of SMU to *F. oxysporum* and *F. solani* hyphae was demonstrated by optical microscopy (c and f, respectively).

We have also performed a time-course dual growth experiment using *F. solani* and SMU for 12 days, which confirmed the results described above, indicating that *F. solani* does not outcompete SMU, even over a longer time period (Figure S1). Importantly, SMU did not display any obvious negative effect on the growth of *Trichoderma* sp. (Figure S1), a well-known plant growth-promoting fungus. This suggests that SMU and *Trichoderma* sp. are compatible and could be tested in combination on inoculant formulations. Finally, the SMU ability to limit *F. solani* growth cannot be merely attributed to the physical occupation of the Petri dish, as a similar effect was not observed when *H. seropedicae*, a well-know PGPR, was used in the dual growth assays (Figure S1).

### SMU substantially increases growth and biomass of maize seedlings

We conducted a pilot gnotobiotic experiment to evaluate whether SMU can promote plant growth, which is the overall effect of the beneficial properties of a PGPR on the host plant. The inoculation of plants can be performed using different methods (e.g. dipping, seed and soil inoculation) (Bashan et al. 2014). We evaluated the potential of SMU in enhancing maize growth *in vivo* by applying a suspension of SMU cells over maize seedlings for 10 days (Figure 4). Our results show that inoculation with SMU led to substantial increases in root and shoot mass (fresh and dry weight), as well as in plant height and radicular length. The biomass increment was 100 % in plant and root length, 80 % for fresh root mass, 64 % for fresh shoot mass and 150 % for dry root and dry shoot mass. Previous studies have shown beneficial effects of other *S. marcescens* isolates on plants, such as in the mitigation of salt stress in wheat (Singh and Jha 2016) and in ginger growth promotion (Dinesh et al. 2015). Another study showed that *S. marcescens* can also be useful in soil phytoremediation (Dong et al. 2014). Thus, our results demonstrate that SMU promote maize growth, probably by a combination of beneficial effects.

**Figure 4:**
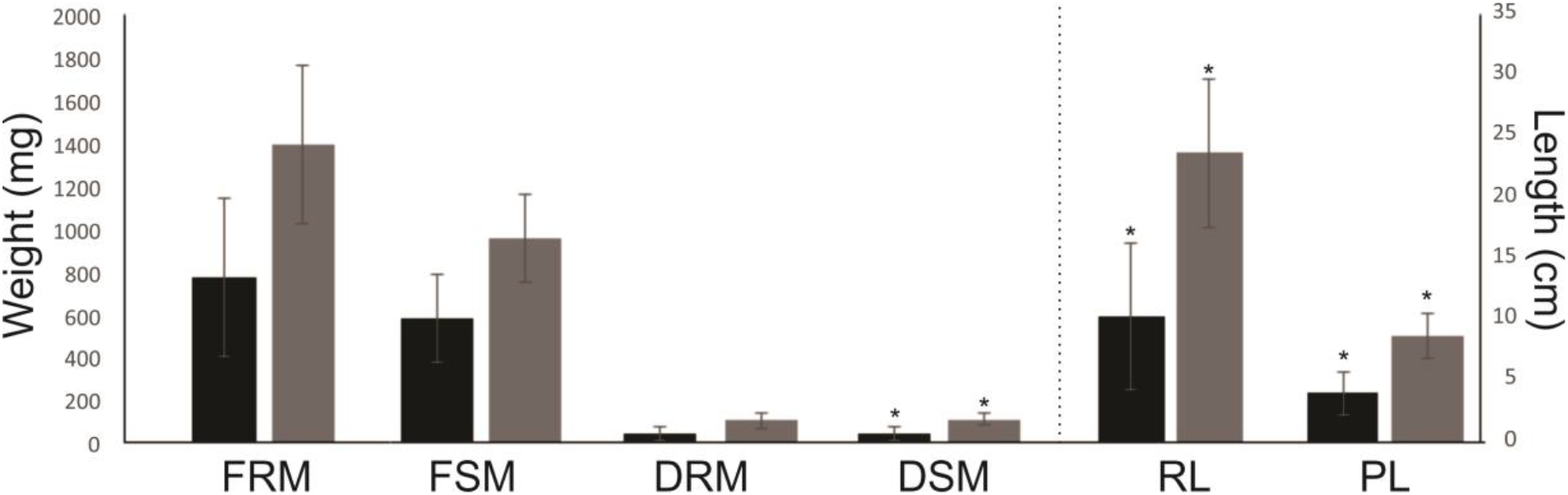
Effect of SMU inoculation on maize seedlings. Germinated seedlings (with 2 to 2.5 cm radicle root length) were transferred to glass tubes containing sterilized vermiculite (one seed per tube). Inoculation was performed by application of 1 mL of the SMU suspension (10^8^ cells⋅mL^−1^) over the seedlings (gray bars). Plants inoculated with 1 mL of the sterile Dygs medium were used as negative controls (black bars). The following metrics were recorded after 10 days: Fresh root mass (FRM), fresh shoot mass (FSM), dry root mass (DRM), dry shoot mass (DSM), root length (RL) and plant height. Experiments were conducted in triplicates and statistical significance assessed by a by Tukey test (p < 0.05).

### Genome structure and comparative analysis

Given the interesting *in vitro* and *in vivo* results, we submitted the SMU genome to whole-genome sequencing using an Illumina Hiseq 2500 instrument (paired-end mode, 2 x 100 bp reads). Sequencing reads were processed with Trimmomatic and assembled with Velvet (see methods for details). The assembled genome consisted of 17 scaffolds (length ≥500 bp) encompassing 5,001,184 bp, with a 59.7 % GC content and an *N*50 of 3,077,593 bp. The genome has 4528 protein-coding genes, 84 and 11 tRNA and rRNA genes, respectively (Figure S2). We used BUSCO (Simao et al. 2015) to estimate genome completeness and detected the complete set of 781 *Enterobacteriales* single-copy genes, supporting the good quality and completeness of the assembled genome (Figure S2). No plasmids were detected in the SMU genome by using plasmidSPADES and Plasmid Finder.

In order to understand genomic features at a species level, we computed the *S. marcescens* pan-genome. A pan-genome is defined as the entire gene repertoire of a given species (Xiao et al. 2015). We used 35 *S. marcescens* isolates with complete or scaffold-level genomes (Table S1). A total of 16,456 gene families were identified, consisting of 2,107 core genes shared by 100 % of the isolates, 7,656 accessory genes shared by more than one and less than 35 isolates and 57 genes unique to SMU (Figure 5a, Table S2). A recent study of 205 clinical strains from the United Kingdom and Ireland reported a pan-genome of 13,614 genes, 3,372 core and 10,215 accessory genes (Moradigaravand et al. 2016). Interestingly, despite the greater number of strains in the clinical study, the reported pan-genome is smaller than that reported here, likely due to the greater diversity of the isolates in our study.

**Figure 5:**
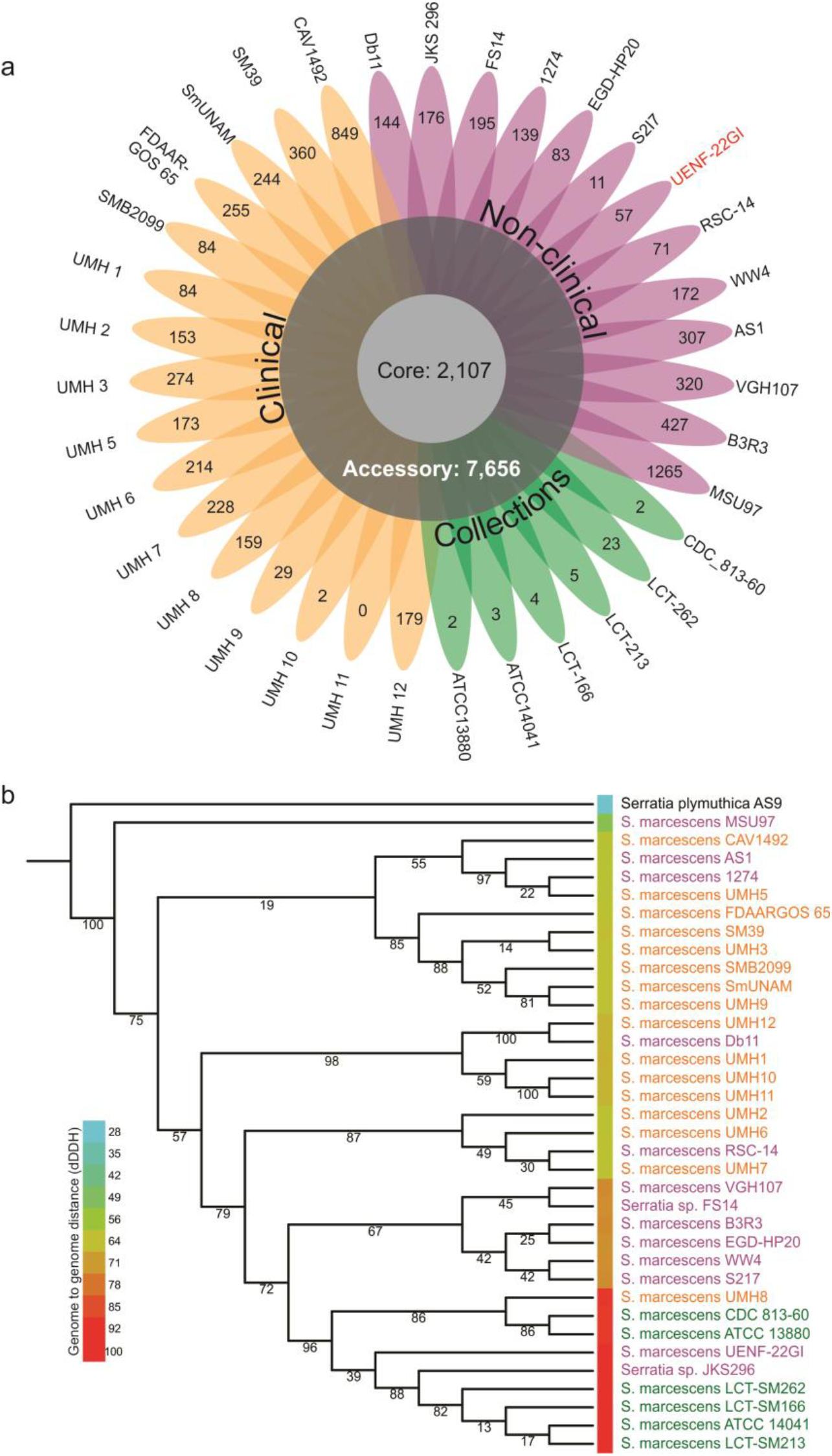
(a) Pan-genome of 35 *S*. marcescens isolates represented as a flowerplot. Clinical, non-clinical and collection *S. marcescens* isolates are represented in yellow, pink and green, respectively. Labels on petal tips represent strain-specific genes. (b) Multi-locus maximum likelihood tree reconstructed using concatenated alignment of 10 single-copy core genes. Branch labels represent bootstrap support (in percentage; 1000 bootstrap replicates). The blue-to-red heatmap accounts for the distance of each isolate to SMU, estimated by the digital DNA:DNA hybridization (dDDH) method.

We performed a phylogenetic reconstruction of the strains included in the pan-genome analysis using 10 single-copy core genes that were also present in the BUSCO reference set. Our phylogenetic reconstructions show a good level of separation of clinical and environmental isolates (Figure 5b). We also computed genome-to-genome distance using the dDDH method (Thompson et al. 2013), which corroborate the structure observed in the phylogenetic tree. Interestingly, a similar partial separation of pathogenic and non-pathogenic strains was also observed in *Burkholderia* and *Paraburkholderia*, respectively (Eberl and Vandamme 2016). Taken together, these results indicate that phylogenetic analysis can also help to assess the applicability of new candidate PGPR. Comparative genomic analysis helped us identify potential genomic islands that might contribute to the competitiveness of this species. In addition to the comparative genomics analysis and phylogenetic reconstructions, we also carefully mined for genes potentially involved in the promotion of plant growth (Table 1). These genes are grouped and discussed according to their general roles, namely: P and Zn solubilization, production of indole compounds (e.g. IAA) and spermidine, biofilm formation, pathogen competition and bioremediation. These findings are discussed in detail in the following sections.

**Table 1:** SMU genes associated with plant-growth promotion features discussed in this study.

| **Phosphate and zinc solubilization** |  |
| --- | --- |
| **Annotation entry (AK961_)** | **Protein name** |
| 07090, 07095, 07100, 07105, 07110 | pqqB, pqqC, pqqD, pqqE, pqqF |
| 10840 | (PQQ)-dependent glucose dehydrogenase |
| 17880 | gluconolactonase |
| 08395 | 2-gluconate dehydrogenase |
| 17580, 17575, 17570 | 2-keto-gluconate dehydrogenase |
| 21125, 21130, 21135, 21140, 21145 | phoU, pstB, pstA, pstC, pstS |
| **Tolerance against metal toxicity** |  |
| 01055 | Arsenate reductase |
| 03990 | ArsR family transcriptional regulator of arsRBC operon |
| 03995 | arsB arsenical pump membrane protein |
| 04000 | arsC1 arsenate reductase |
| 01905, 01915 | Copper resistance protein |
| 09290 | Copper resistance protein copD |
| 07435, 07440 | Chromate transporter |
| 12615 | Cobalt-zinc-cadmium efflux system |
| **IAA and spermidine-related** |  |
| 01575 | ipdC |
| 00655, 12310 | Auxin efflux carrier |
| 18130, 18125 | speAB |
| 18275, 18270 | speDE |
| **Biofilm formation** |  |
| 20475, 20470, 20465, 20460, | bcsA, bcsB, bcsC, bcsZ |
| 20480, 20485 | bcsQ, bcsR |
| 20490, 20495, 20500 | bcsE, bcsF, bcsG |
| 01650, 01655, 1660, 01665 | pgaA, pgaB, pgaC, pgaD |
| 13115 | adrA |
| **Biocontrol and resistance** |  |
| 20530, 01270, 12935, 05475 | chiA, chiB, chiD, chiA1 |
| 13300, 13305, 13310, 13315, | pigA, pigB, pigC, pigD, pigE, pigF, pigG, pigH, pigI, pigJ, pigK, pigL, |
| 13320, 13325, 13330, 13335, | pigM, pigN |
| 13340, 13345, 13350, 13355, |  |
| 13360, 13365 |  |
| 16235 | Kasugamycin resistance protein ksgA |
| 03740, 14040 | Bicyclomycin resistance protein |
| 02590 | Bicyclomycin multidrug efflux system |
| 13395 | Fosmidomycin resistance protein |
| 07380 | Barnase inhibitor |
| 08005 | Fusaric acid resistance protein |

### Identification and analysis of the horizontally transferred Gap1 island

Our comparative analysis uncovered a remarkable region in the SMU genome that is absent in most other *S. marcescens* genomes, which we named as Gap1 (Figure 6a). However, Gap1 is partially conserved in the JSK296 and ATCC14041 strains (Figure 6b), which belong to the SMU phylogenetic clade. Although partially eroded in several members of the clade, this result lends additional support to the greater proximity of SMU to a group of environmental strains (Figure 6b). Manual analysis assisted by results from IslandViewer allowed us to predict that this ~52 Kb long genomic island contains 38 genes. This island encodes its own integrase of the tyrosine recombinase superfamily (AK961_03610), which is also encoded by several phages and bacterial mobile elements (Iyer and Aravind 2012), suggesting that it supports its own genetic mobility. Genomic islands with closely related genes were also detected in several distantly related proteobacteria, such as *Erwinia piriflorinigrans* CFBP 5888, *Erwinia sp*. ErVv1, *Hahella sp*. CCB-MM4, *Enterobacter sp*. T1-1 and *[Polyangium] brachysporum*, suggesting that fitness-conferring determinants carried by this island might have facilitated their dissemination by HGT. Further, the closest cognates of at least 15 genes (AK961_03495: AK961_03565) in this island are found in *Erwinia* species raising the possibility of a relatively recent genetic exchange event involving *Serratia* and *Erwinia*.

**Figure 6:**
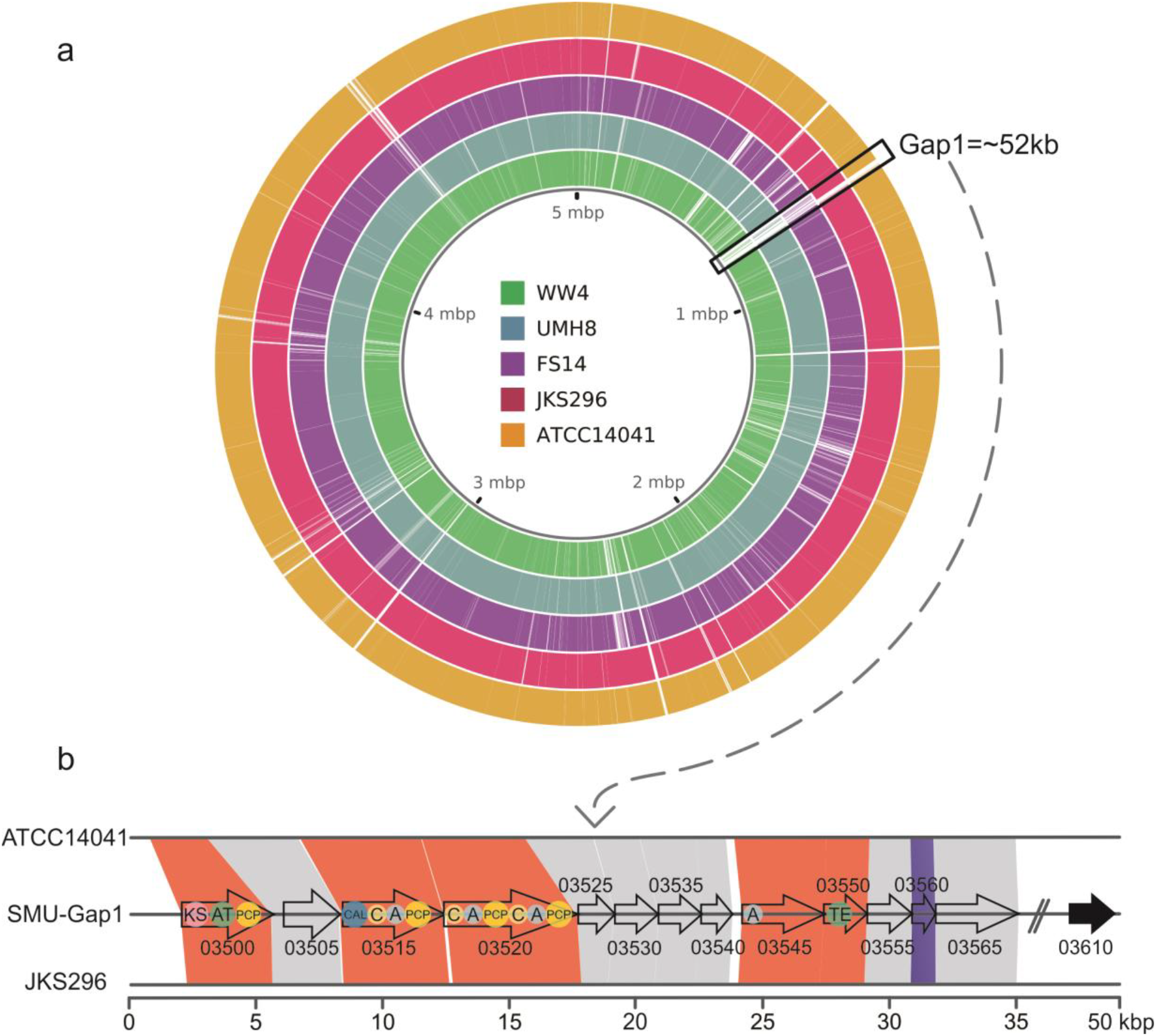
(a) Whole-genome alignment of SMU and some of the closest reference genomes. The black box indicates the horizontally-acquired region (Gap1); (b) Synteny analysis of part of the genes within Gap1 region, emphasizing the presence of the NRPS-PKS domains: KS (ketosynthase), AT (acyltransferase), PCP (peptidyl carrier domain), CAL (coenzyme A ligase), C (condensation), A (adenylation) and TE (thioesterase). AK961_03560 encodes an antitoxin protein. AK961_03610 encodes an integrase that likely delimits the end of Gap1.

We next investigated the island to identify genes potentially functioning as fitness determinants which could explain this wide dissemination. At the 5’ flank of the island are two genes respectively encoding a JAB domain protein of the RadC family and ArdB domain (AK961_03485, AK961_03490). These genes were recently identified as part of a system of proteins that enable mobile elements such as conjugative transposons and plasmids to evade restriction by host defense systems (Iyer et al. 2017). The core of the island contains an operon, which is shared with the related islands that we detected in the above-stated bacteria, encoding the system predicted to synthesize a non-ribosomal peptide. The two largest genes (AK961_03515, AK961_03520) of this operon code for two giant multidomain non-ribosomal-peptide synthetases (NRPS), together with 4 predicted AMPylating domains that charge acyl groups and 3 condensation domains that ligate charged amino acids to form a peptide bond. Additionally, the operon contains a further gene for a standalone AMPylating enzyme and one for a thioesterase of the α/β-hydrolase fold (AK961_03545, AK961_03550). The last enzyme has been shown to be required for generation of a cyclic peptide in several NRPS systems (Schneider and Marahiel 1998). Thus, the system encoded by this island has the potential to synthesize tetra- or penta-peptide skeleton with a possibly cyclic structure. Notably, the region also encodes a GNAT acetyltransferase (AK961_03495) that might either modify this peptide or confer auto-resistance against its toxicity. Also in this operon is a gene for a pol-β superfamily nucleotidytransferase (AK961_03530), which might modify the peptide generated by the NRPS by the addition of a nucleotide, or regulate its production/secretion by nucleotidylation of one of the components of the system. The said operon codes for a predicted peptide transporter of the MFS superfamily that probably facilitates the export of the synthesized peptide out of the cell. Taken together, we interpret this NRPS system and associated proteins are generating an anti-microbial peptide.

We also found this island to encode a protein belonging to a previously unknown family of the ADP-ribosyltransferase (ART) fold (AK961_03540) (Aravind et al. 2015). Using sequence profile searches and profile-profile comparisons we showed that this novel family also includes the Pfam (“Domain of unknown function”) DUF4433 and the abortive phage infection protein AbiGi. Members of the ART superfamily utilize NAD^+^ to either transfer it to target substrates (e.g. proteins) or degrade NAD^+^. Given the relationship to the AbiGi proteins, we predict that this protein might also play a role in anti-phage defense by means of its ADP-ribosyltransferase activity targeted either at self or viral proteins. In a similar vein, we also found an ATPase of the ABC superfamily (AK961_03575) that is related to AbiEii, another abortive infection protein involved in anti-phage systems. More remarkably, the gene encoding this protein is also part of an operon coding for a KAP NTPase (AK961_03580) and another protein (AK961_03585), which are a version of an anti-phage system centered on these two proteins (Aravind et al. 2004). Of these the proteins, AK961_03585 is predicted to function as a novel DNA transglycosylase that is predicted to incorporate a modified base into DNA, which is likely to be a deazaguanine acquired from the queuine biosynthesis pathway (Iyer et al. 2013).

Taken together, this island codes for multiple distinct fitness-promoting systems: one predicted to synthesize a potential antimicrobial peptide that could be deployed against competing organisms in compost. Further, it also encodes a beta-lactam amidase (AK961_03565) that could likely defend SMU against certain beta-lactams (e.g. penicillin) produced by competing bacteria. The further set of genes is likely to confer resistance against certain bacteriophages and potentially enhance the fitness of this strain relative to other *Serratia* lacking the island.

### Phosphorus and zinc solubilization genes

As discussed above, in tropical environments, P is mostly present in poorly soluble mineral phosphates that are not readily available for plant uptake (An and Moe 2016). Microbial conversion of insoluble mineral P forms into soluble ionic phosphate (H_2_PO_4_^−^) is a key mechanism of increasing the P availability (Alori et al. 2017). Further, the production and secretion of a variety of low molecular weight acids constitute a major strategy to solubilize not only P (An and Moe 2016), but also Zn (Solanki et al. 2016). Among these substances, gluconic acid, produced by three oxidation reactions carried out by membrane-bound periplasmic proteins (Krishnaraj and Goldstein 2001), is typically the most prominent.

The SMU genome harbors a number of genes involved in the production of gluconic acid from glucose (Table 1), which starts with the oxidation of glucose by a membrane-bound, periplasmic pyrroloquinoline-quinone (PQQ)-dependent glucose dehydrogenase (GDH; AK961_10840). The intermediate glucono-1,5-lactone is hydrolyzed to gluconate by a gluconolactonase (AK961_17880) and oxidized by 2-gluconate dehydrogenase (AK961_08395) to 2-ketogluconate, which is oxidized to 2-5-diketo gluconate by 2-keto-gluconate dehydrogenase, an enzymatic complex comprising a small (AK961_17580), a large (AK961_17575) and a cytochrome (AK961_17570) subunits (as in *Gluconobacter oxydans*, accession AB985494), encoded in the same operon. Gluconic acid synthesis requires the PQQ cofactor (Duine 1991), which is produced by proteins encoded by the *pqqBCDEF* operon. Importantly, this operon is fully conserved in the SMU genome (genes AK961_07090: AK961_07110) (Table 1; Figure S3). The SMU genome also has a conserved *pstABCS* operon (AK961_21130: AK961_21145) (Table 1; Figure S2b), which encodes a phosphate-specific transport system. Finally, current data indicate that Zn solubilization is largely carried out by the same genes involved in the solubilization of inorganic P (Intorne et al. 2009). Therefore, based on genomic data and *in vitro* evidence, we hypothesize that SMU solubilizes P and Zn through soil acidification.

### Tolerance against metal toxicity

Successful soil bacteria often have to tolerate metal contamination, which can involve different strategies (Das et al. 2016). In addition, PGPR can alleviate the impact of heavy metals on plants by reduction, oxidation, methylation and conversion to less toxic forms (Hassan et al. 2017). We found a number of genes related to these roles in the SMU genome (Table 1): arsenate reductase (AK961_01055), *arsRBC* (AK961_03990, AK961_03995, AK961_04000), copper resistance protein (AK961_01905, AK961_01915, AK961_09290), *cusRS* (AK961_11430, AK961_11425), chromate transporter ChrA (AK961_07435, AK961_07440) chromate reductase (AK961_21085) and *czcD* (AK961_12615). Although this list is likely incomplete due to the wide diversity of reactions and pathways involved in these tolerance pathways, our findings are in line with those from a recently sequenced genome of a *S. marcescens* strain that alleviates cadmium stress in plants (Khan et al. 2017). Notably, a gene from the Gap1 island (AK961_03595) codes for a member of the YfeE-like transporter family, which transport chelated Fe/Mn and could potentially play a role in alleviating toxicity from these transition metals.

### IAA, spermidine biosynthesis and phenolic compound transport

We searched for IAA biosynthesis pathways in the SMU genome and found the *ipdC* gene (Table 1). This gene encodes a key enzyme responsible for the conversion of indole-3-pyruvate in indole-3-acetaldehyde, a critical step of the IPyA pathway. Disruption of *ipdC* dramatically decreases IAA production in *A. brasilense* (Malhotra and Srivastava 2008). In addition, we have also identified two putative auxin efflux carrier genes (AK961_00655, AK961_12310) (Table 1), suggesting that SMU also exports IAA. These results indicate that the IPyA pathway is active in SMU. Other IAA biosynthesis pathways were only partially identified and the genes pertaining to these pathways are also part of other processes. Hence, activity of alternative pathways in SMU warrants further investigation.

In addition to IAA, we have also found the *speAB* (AK961_18130, AK961_18125) and *speDE* (AK961_18275, AK961_18270) operons, which are involved in spermidine biosynthesis (Table 1). Polyamines (e.g. spermidines) are essential for eukaryotic cells viability and have been correlated with lateral root development, pathogen resistance and alleviation of oxidative, osmotic and acidic stresses (Xie et al. 2014). Therefore, spermidine production by SMU may constitute an additional mechanism involved in plant-growth promotion.

Several plants produce phenolic compounds, which are part of their defense system and are also regulators of their own growth. Interestingly, the Gap1 island codes for a 4-hydroxybenzoate transporter (AK961_03620), which is closely related to cognate transporters from other plant-associated bacteria, such as *Pantoea ananatis*, *Erwinia amylovorans*, *Pseudomonas putida*, and *Dickeya* species. This suggests that this transporter might play a role in the plant-bacterium interaction via phenolic compounds such as benzoate, as has been proposed for certain *Xanthomonas* species (Wang et al. 2015).

### Biofilm formation and biocontrol of phytopathogenic fungi

Bacterial biofilms are multicellular communities entrapped within an extracellular polymeric matrix (Flemming et al. 2016) that are essential for survival, microbe-microbe interactions and root colonization (Kasim et al. 2016). We found several biofilm related genes in the SMU genome (Table 1), such as *pgaABCD* (AK961_01650: AK961_01665) (Figure S3). This operon is responsible for the production of poly-β-1,6-N-acetyl-D-glucosamine (PGA), which is associated with surface attachment, intercellular adhesion and biofilm formation in several species (Echeverz et al. 2017).

Cellulose is the fundamental component of plant cell walls and the most abundant biopolymer in nature. Cellulose biosynthesis has been also described in a broad range of bacteria and a variety of bacterial cellulose synthase operons are known (Römling and Galperin 2015). In proteobacteria, cellulose biosynthesis is mainly carried out by the *bcsABZC* and *bcsEFG* operons, along with the *bcsQ* and *bcsR* genes, described as the *E. coli*-like *bcs* operon (Krasteva et al. 2017). The *bcsABZC* (AK961_20475, AK961_20470, AK961_20465, AK961_20460) and *bcsEFG* (AK961_20490, AK961_20495, AK961_20500) operons are proximal to each other in the SMU genome, although in opposite strands (Figure S3). The opposite orientation of these operons is also observed in others *S. marcescens* strains (e.g. WW4, B3R3 and UMH8), and might be related to the transcriptional regulation of biofilm synthesis in *S. marcescens*. Further, there are two regulatory genes upstream to the *bcsABZC* operon: *bcsQ* (AK961_20480) and *bcsR* (AK961_20485). These regulatory genes were also reported to be required for cellulose synthesis and subcellular localization of an active biosynthesis apparatus at the bacterial cell pole in γ-proteobacteria (Le Quéré and Ghigo 2009). We have also found the *adrA* gene (AK961_13115), which encodes a diguanylate cyclase that synthesizes cyclic dimeric GMP, which binds to the BcsA and activates cellulose production (Cowles et al. 2016). BcsA has two cytoplasmic domains and transmembrane segments, while BcsB is located in the periplasm, anchored to the membrane; together they form the BcsAB complex, which function as a channel for the addition of new residues to the nascent glucan molecule (Römling and Galperin 2015). BcsC is an outer membrane pore (Whitney and Howell 2013) and BcsZ is an endoglucanase that may be involved in the alignment of β-glucans prior to export (Castiblanco and Sundin 2016), or act as negative regulator of cellulose production (Ahmad et al. 2016). The *bcsEFG* operon is also necessary for optimal cellulose synthesis (Fang et al. 2014) and its deletion disrupted cellulose production (Serra et al. 2013).

### Fungi biocontrol, prodigiosin production and resistance to antimicrobial compounds

Chitinases are central to the catabolism of chitin (i.e. poly β-(1->4)-N-acetyl-D-glucosamine), constituting a route by which bacteria can access a rich source of nutrients (Paspaliari et al. 2017). Chitinases break chitin into soluble oligosaccharides that can be transported into the periplasm via a chitoporin channel, where they are further processed into mono- and di-saccharides that are transported to the cytoplasm (Hayes et al. 2017). Because of the severe impact of phytopathogenic fungi in agriculture (Santamarina et al. 2017), chitinases have received increased attention by the scientific community as a biocontrol mechanism deployed by several bacteria, including *S. marcescens* (Vaikuntapu et al. 2016). We found 4 chitinases in the SMU genome (Table 1); to further classify them we performed BLASTp searches on the Swissprot database. AK961_20530 shares 99 % identity with chitinase A (accession: P07254), AK961_01270 shares 100 % identity with chitinase B (accession: P11797), AK961_12935 shares 29 % identity and 93 % coverage with chitinase D (accession: P27050) and AK961_05475 shares 31 % identity and 88 % coverage with chitinase A1 (accession: P20533). We have also found other chitin metabolism genes in SMU, namely AK961_01260 and AK961_12890, which encode a chitin-binding protein and a chitobiase, respectively. Bacterial chitinases have been reported to compromise fungal spore integrity and generate germ tube abnormalities (Pandey et al. 2016). Further, ChiA promotes the degradation of mycelia of several phytopathogenic fungi, including *Fusarium, Acremonium* and *Alternaria* species (Medina-de la Rosa et al. 2016). Chitinase applications are not restricted to fungal biocontrol and can also be deployed for bioremediation and bioconversion of chitin wastes, as well as part of an insect biocontrol strategies (Hamid et al. 2013).

As part of the arms race between microorganisms, several *Fusarium* species produce fusaric acid, a mycotoxin reported to be toxic to some microorganisms, such as *P. fluorescens* (Crutcher et al. 2017). Interestingly, SMU has a gene that encodes a fusaric acid resistance protein (AK961_08005), which may render SMU resistant to this toxin, indirectly boosting its fungicidal activity.

We have also found a widely-conserved operon comprising genes involved in the biosynthesis of prodigiosin (i.e. the *pig* operon), the notorious red pigment observed in several *Serratia* isolates (Mahlen 2011). The *pig* operon in the SMU genome comprises 14 genes (Table 1, Figure S3) arranged in a structure that resembles the *pig* operon from *S. marcescens* ATCC274 (also an environmental isolate) (Harris et al. 2004). Prodigiosin is most commonly found in environmental *S. marcescens* isolates and has been proposed to suppress growth of various fungi (Duzhak et al. 2012), bacteria (Danevčič et al. 2016), protozoans (Genes et al. 2011) and even viruses (Zhou et al. 2016). It has been recently suggested that prodigiosin has affinity for the lipid bilayer of the plasmatic membrane, causing outer membrane damage (Darshan and Manonmani 2016). Although the mechanistic details of the prodigiosin antifungal and antibacterial activities remain largely unclear, our findings on the dual growth experiments indicate that that the *pig* operon is active in SMU and that prodigiosin delineates the growth area of *F. solani* (Figure 3, Figure S1).

*Streptomyces* species are ubiquitous in the soil and notable for the production of several antimicrobials (Barka et al. 2016). Therefore, the presence of genes conferring resistance against these antimicrobials is a desirable feature of a successful PGPR. SMU produces several resistance proteins against *Streptomyces* antimicrobials such as bicyclomycin (AK961_03740, AK961_14040, AK961_02590), fosmidomycin (AK961_13395) and kasugamycin (AK961_16235). In addition, SMU also exhibits a type 6 secretion system (T6SS) (AK961_04125-AK961_04210) that can mediate interbacterial antagonistic interactions (Russell et al. 2014; Zhang et al. 2012).

## CONCLUDING REMARKS

In this study we assessed the plant growth-promoting properties of SMU using *in vitro* biochemical assays and *in vivo* experiments in greenhouse conditions. Specifically, we found that SMU is able to: 1) solubilize inorganic P and Zn; 2) produce indole compounds; 3) counter the growth of two phytopathogenic *Fusarium* species by a combination of physical (i.e. biofilm formation) and biochemical (e.g. prodigiosin, chitinase) properties and; 4) substantially increase growth and biomass of maize seedlings. Given these interesting properties, we sequenced the SMU genome and mapped the genes that are likely responsible for these traits. Interestingly, the SMU genome also harbors a mobile genomic island comprising 38 genes that were horizontally transferred. This region codes for a NRPS systems and other proteins predicted to confer fitness advantage by various mechanisms, including DNA modification and anti-phage defenses. Phylogenetic analysis show that SMU groups within a clade comprised almost exclusively of non-clinical isolates. Together with the absence of plasmids and synthesis of prodigiosin, more frequent in environmental isolates, we hypothesize that SMU is non-pathogenic. However, basic safety issues must be addressed before biotechnological applications can be envisaged. Collectively, our results add important information regarding *S. marcescens* plant growth-promoting abilities that can inspire future applications in inoculant formulations.

## MATERIALS AND METHODS

### Vermicompost maturation

Mature vermicompost was produced with dry cattle manure as substrate inside a 150 L cement ring. Humidity was kept at 60-70%, by weekly watering and mixing. After 1 month, earthworms (*Eisenia foetida*) were introduced at the rate of 5 kg⋅m^3^. After 4 months, earthworms were removed and the vermicompost was placed in plastic bags and stored at 25°. At the final maturation stage, the chemical composition of the substrate (in g⋅kg^−1^) was as follows: total nitrogen (1.9 ± 0.4); total carbon (22.99 ± 3.3); P_2_O_5_ (6.97 ± 1.4); C/N ratio of 13.8 ± 0.4 and pH (H_2_O) = 6.6 ± 0.18.

### Bacterial isolation and DNA purification

Serial dilutions were performed on a solution prepared by adding 10 g of vermicompost in 90 mL of saline (8.5 g⋅L^−1^ NaCl), followed by shaking for 60 minutes. Next, 1 mL of the initial dilution (10^−1^) was added to a new tube containing 9 mL of saline (10^−2^), and successively until 10^−7^ dilution. Then, 100 μL of the final dilutions from 10^−5^ to 10^−7^ were taken and spread on plates containing solid Nutrient Broth (NB) with 8 g⋅L^−1^ of NB and 15 g⋅L^−1^ of agar in 1 L of distilled water. After incubation at 30 °C for 7 days, different colony types could be identified and, for purification, individual colonies were transferred to Petri plates with Dygs solid media (D bereiner et al. 1995) containing 2 g⋅L^−1^ of glucose, 2 g⋅L^−1^ of malic acid, 1.5 g⋅L^−1^ of bacteriological peptone, 2 g⋅L^−1^ of yeast extract, 0.5 g⋅L^−1^ of K_2_HPO_4_, 0.5 g⋅L^−1^ of MgSO_4_⋅7H_2_O, 1.5 g⋅L^−1^ of glutamic acid and 15 g⋅L^−1^ of agar, adjusted to pH 6.0; these supplies were acquired from Vetec (São Paulo, Brazil). From the last dilution (10^−7^) and after the isolation and purification on Dygs solid medium, a pink-to-red, circular, pulvinate elevation, punctiform and smooth surface bacterial colony was selected. Phase contrast microscopy revealed the presence of Gram-negative, rod-shaped and non-motile cells. This distinctive isolate, named UENF-22GI, was stored in 16 mL glass flask containing 5 mL of Nutrient Broth solid medium covered with mineral oil and later grown in liquid Dygs medium under rotatory shaker at 150 rpm and 30 °C for 36 h to perform *in vitro* and *in vivo* assays. Total DNA of UENF-22GI was extracted using QIAamp® DNA Mini Kit (QIAGEN GmbH, Hilden, Germany). DNA quantification and quality assessment were performed using an Agilent Bioanalyzer 2100 instrument (Agilent, California, USA).

### Phosphorus and zinc solubilization

Bacterial inocula were grown for 36 h on liquid Dygs media at 150 rpm and 30 °C until approximately 10^8^ cells.mL^−1^ (O.D._540nm_ = 1.0) (D bereiner et al. 1995). To carry out a qualitative P solubilization assay, 10 µl of the bacterial suspension were added to petri dishes containing 10 g⋅L^−1^ of glucose, 5 g⋅L^−1^ of ammonium chloride (NH_4_Cl), 1 g⋅L^−1^ of sodium chloride, 1 g⋅L^−1^ of magnesium sulfate heptahydrate (MgSO_4_⋅7H_2_O), 15 g⋅L^−1^ of agar in 1 L of distilled water at pH 7.0, and incubated at 30 °C for 7 days. Two mineral P sources were tested: calcium phosphate Ca_3_(PO_4_)_2_ (P-Ca) and fluorapatite rock phosphate Ca_10_(PO_4_)_6_F_2_ (P-rock), both at 1 g⋅L^−1^. Positive P solubilization phenotypes were based on halo formation around bacterial colonies and results were expressed in the form of a Solubilization Index (SI), calculated as the halo diameter (d1) divided by the colony diameter (d2). The SI values can be used to classify the solubilization ability of a strain as low (SI < 2), intermediate (2 < SI < 4) and high (SI > 4) (Marra et al. 2015).

Quantitative P solubilization assays were also performed. 50 μL bacterial suspensions in Dygs liquid medium were transferred to 30 mL test tubes containing Pikovskaya liquid medium at pH 7.0, supplemented with P-Ca or P-rock at 1 g⋅L^−1^. The assay was carried out in orbital shaker at 150 rpm at 30 ºC. After 7 days growth, a 5 mL aliquot was harvested and centrifuged at 3200 rpm for 15 min. The supernatant was used to determine the pH and to quantify soluble P levels by the colorimetric ammonium molybdate method (λ = 600 nm). Results were expressed in mg of PO ^2-^⋅L^−1^.

Zn solubilization was evaluated using 10 µL aliquots taken from the bacterial suspension and dropped onto petri dishes containing solid media (Saravanan et al. 2007) constituted of 10 g⋅L^−1^ of glucose, 1 g⋅L^−1^ of ammonium sulfate ((NH_4_)_2_SO_4_), 0.2 g⋅L^−1^ of potassium chloride (KCl), 0.1 g⋅L^−1^ of dipotassium phosphate (K_2_HPO_4_), 0.2 g⋅L^−1^ of magnesium sulfate heptahydrate, 1.0 g⋅L^−1^ of zinc oxide (ZnO), 15 g⋅L^−1^ agar, 1 L distilled water; the medium was incubated for 7 days at 30 °C. Zn solubilization was also assessed by halo formation around bacterial colonies. Both, Zn and P solubilization assays were carried out in triplicates.

### Production of indole compounds

To quantify the production of indole compounds, previously grown bacteria were transferred to glass tubes containing 5 mL of Dygs medium with or without tryptophan addition (100 mg⋅L^−1^), followed by 72 h incubation in the dark, at 30 °C and 150 rpm. To evaluate indole synthesis (Sarwar and Kremer 1995), 150 µL of grown bacteria were transferred to microplates and 100 µL of Salkowski reagent, which was prepared by diluting 1 mL of an iron trichloride hexahydrated (FeCl_3_⋅6H_2_0) aqueous solution at 92.5 g⋅L^−1^ in 50 mL of perchloric acid (HClO_4_) 350 g⋅L^−1^ in water. The plate was incubated for 30 min in the dark and samples analyzed at 492 nm on a UV mini 1240 spectrophotometer (Shimadzu, Japan). This assay was conducted in triplicate.

### *In vitro* dual culture assays

*In vitro* bacterial-fungal dual culture assays were performed in 9 cm diameter Petri dishes containing Potato Dextrose Agar solid medium. A 5 mm diameter disk taken from the edge of actively growing hyphae of *F. solani* and *F. oxysporum* were inoculated at the center of each Petri dish. Suspensions of SMU were spotted in four equidistant quadrant points to the inoculated fungal disk. Control treatments (without SMU) were conducted in parallel to monitor fungal growth. Treatments were carried out for 10 days and three independent replicates were performed. In addition, time-course dual culture experiments were also performed for 12 days with SMU and *F. solani* or *Trichoderma* sp., a plant growth-promoting fungus. We have also tested *F. solani* in dual growth assays with *H. seropedicae* HRC54, a well-known PGPR without known anti-fungal properties. Samples from the transition zones between fungi structures and spotted bacteria were mounted on glass slide and coverslip, observed under phase-contrast inverted optical microscope Zeiss Axio 10 Observer A1 and photodocumented with an Axiocam MRC 5 digital camera. For the time-course assays between *F. solani* and SMU (1 −12 days of growth), bacterial inocula were spotted in three equidistant quadrant points to the inoculated fungal disk.

### *In vivo* plant-growth promotion assays

Maize (*Zea mays* var. UENF/506-11) seeds were surface-disinfected using ethanol 70 % for 30 seconds, followed by a wash with 5 % sodium hypochlorite (NaClO) for 20 min. Next, seeds were washed 5 times with sterile distilled water under stirring for 3 minutes and transferred to petri dishes containing 1.5 % solidified agar for pre-germination for 4 days. Seedlings with 2.0 to 2.5 cm radicle length were carefully transferred under flow chamber to glass tubes of 2 cm diameter and 20 cm height containing 10 g of sterilized vermiculite (one seed per tube). Meanwhile, the bacterial inoculum was prepared by growth in Dygs liquid media for 36 h, at 30 °C and 120 rpm. Inoculation was performed by application of 1 mL of the SMU suspension (10^8^ cells⋅mL^−1^) over the seedlings. Plants inoculated with 1 mL of sterile Dygs medium were used as negative controls. The assay was carried out under laboratory conditions with average temperature at 30 °C and 12 h of light/dark photoperiod. After 10 days, plants were collected and the following biometric measurements were registered: height (cm), total radicular length (cm), fresh root mass (mg), fresh shoot mass (mg), dry root mass (mg) and dry shoot mass (mg). This assay was performed in four replicates. Statistical analyses were performed using the SAEG software (Universidade Federal de Viçosa, Brazil) and obtained means were compared with the Tukey test.

### Genome sequencing and assembly

Paired-end libraries were prepared with the TruSeq Nano DNA LT Library Prep (Illumina) and sequenced on a HiSeq 2500 instrument at the Life Sciences Core Facility (LaCTAD; UNICAMP, Campinas, Brazil). The quality of the sequencing reads (2 x 100bp) was checked with FastQC 0.11.5 (https://www.bioinformatics.babraham.ac.uk/projects/fastqc/). Quality filtering was performed with Trimmomatic 0.35 (Bolger et al. 2014) and only reads with average quality greater than 30 were used. The UENF-22GI genome was assembled with Velvet 1.2.10 (Zerbino and Birney 2008), with the aid of VelvetOptimiser 2.2.6 (Gladman and Seemann 2008). Scaffolding was performed with SSPACE 3.0 with default parameters (Boetzer et al. 2011). QUAST 4.0 (Gurevich et al. 2013) was used to assess general assembly statistics. Genome completeness was assessed with BUSCO 3.0 (Simao et al. 2015), using the *Enterobacteriales* dataset as reference.

### Genome annotation and phylogenetic analysis

The assembled genome was annotated with the NCBI Prokaryotic Genome Annotation Pipeline (Tatusova et al. 2016). Some annotations were manually improved with primary literature information and specific searches using BLAST (Altschul et al. 1997) and Kegg Orthology And Links Annotation (BlastKOALA) (Kanehisa et al. 2016). The presence of plasmids was assessed with plasmidSPAdes 3.10 (Antipov et al. 2016) and PlasmidFinder 1.3 (Carattoli et al. 2014). Genes and operons involved in antibiotic and secondary metabolism were predicted using antiSMASH 4.0 (Blin et al. 2017). The SMU genome was deposited on Genbank under the BioProject PRJNA290503.

Whole genome comparisons were done using BRIG 0.95 (Alikhan et al. 2011) and synteny was assessed using Synima v 1.0 (Farrer 2017). Horizontal gene transfer regions were inferred with IslandViewer4 (Bertelli et al. 2017) followed by manual adjustments. Pan-genome analysis was performed with BPGA 1.3.0 (Chaudhari et al. 2016). Phylogenetic reconstructions were carried out using the predicted proteins of 10 core housekeeping genes that were also present in the BUSCO’s reference dataset. Protein sequences were aligned using MUSCLE 3.8.31 (Edgar 2004) and evolutionary model selected with protest 3.4.2 (Darriba et al. 2011). Maximum-likelihood phylogenetic reconstructions were performed using RAxML 8.2.10 (Stamatakis 2014), with the Le and Gascuel model (Le and Gascuel 2008), gamma correction, SH local support and 1000 bootstrap replicates. Genomic distance patterns were computed with the digital DNA:DNA hybridization (dDDH) (Auch et al. 2010). The resulting phylogenetic tree and dDDH values were integrated and rendered in iTOL 3 (Letunic and Bork 2016).

## ACKNOWLEDGEMENTS

This work was supported by Fundação Carlos Chagas Filho de Amparo à Pesquisa do Estado do Rio de Janeiro, Conselho Nacional de Desenvolvimento Científico e Tecnológico (CNPq) and Coordenação de Aperfeiçoamento de Pessoal de Nível Superior (CAPES). We would like to thank the staff of the Life Sciences Core Facility (LaCTAD) of UNICAMP for library preparation and genome sequencing. We also thank Prof. Valdirene Gomes (UENF) for kindly providing the *Fusarium* strains and to Francisnei Pedrosa for helping in figure preparation.

