## Supplementary Materials for "Genome sequencing and assessment of plant growth-promoting properties of a *Serratia marcescens* strain isolated from vermicompost"

[a] Laboratório de Química e Função de Proteínas e Peptídeos, Universidade Estadual do Norte Fluminense Darcy Ribeiro (UENF), Rio de Janeiro, Brazil; [b] Núcleo de Desenvolvimento de Insumos Biológicos para a Agricultura (NUDIBA), Universidade Estadual do Norte Fluminense Darcy Ribeiro (UENF), Rio de Janeiro, Brazil. [c] Departamento de Bioquímica, Universidade Federal do Paraná (UFPR), Paraná, Brazil; [d] National Center for Biotechnology Information, National Library of Medicine, National Institutes of Health, Bethesda, Maryland, United States of America.

[#] Corresponding authors:

Fabio L. Olivares:

Thiago M. Venancio:

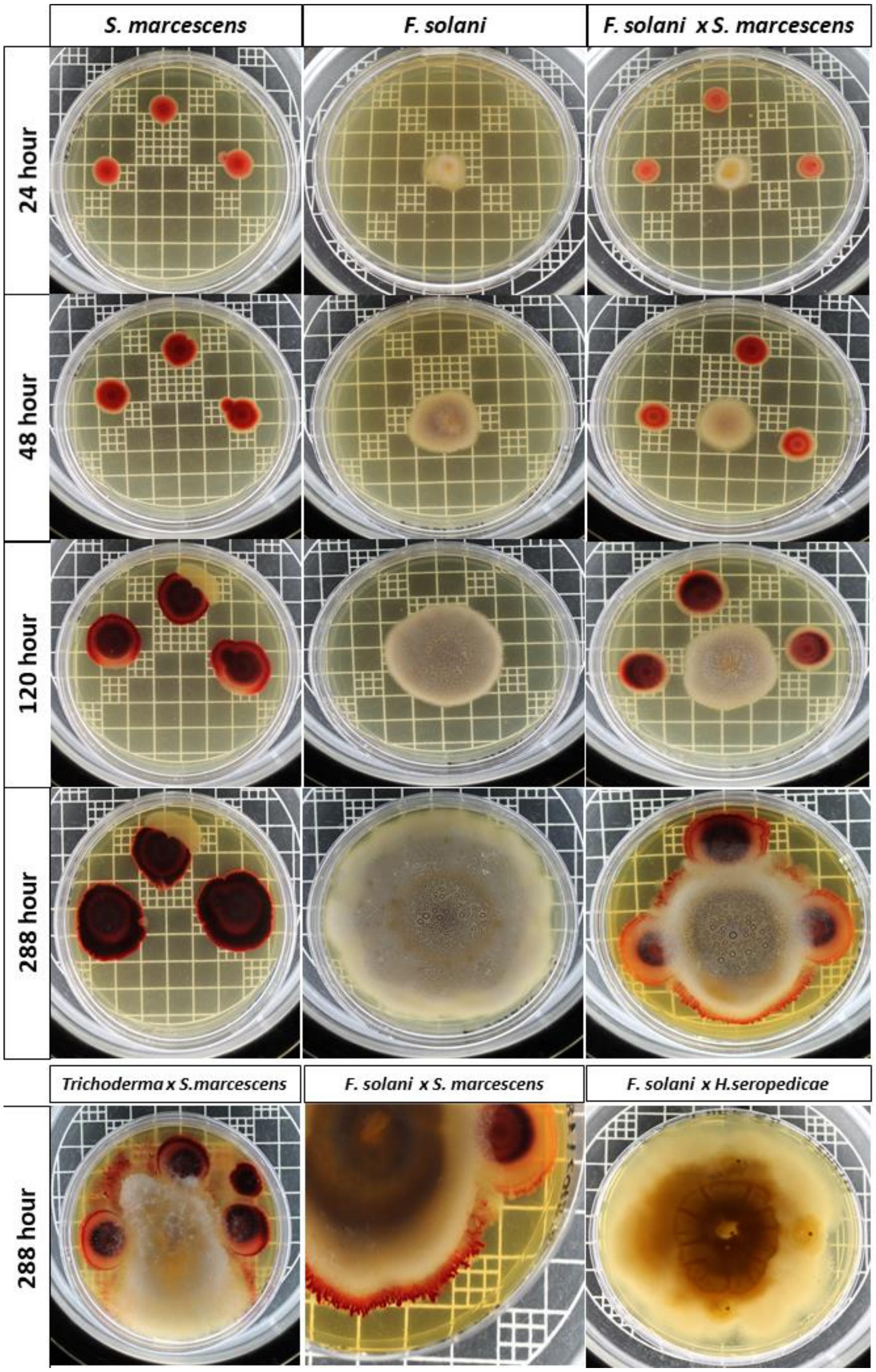


**Figure S1:** Time-course (24, 48, 120 and 288 h) dual growth of SMU with the phytopathogenic *Fusarium solani* on potato dextrose agar (PDA) solid medium. Note that *F. solani* colony growth (i.e. spread) was reduced by SMU, which in contrast was not significantly affected by the fungus, despite some alterations in the pigmentation patterns. At the bottom line, we show that SMU does not counter the growth of the beneficial saprophytic fungus *Trichoderma* sp.. We also observed a depigmentation of the SMU colony and its spread on the plate surrounding *Trichoderma* sp.. Finally, we used another bacteria species, *Herbaspirillum seropedicae*, to demonstrate that the SMU effects on *Fusarium* are not spurious or merely due to physical occupation of the Petri dish.

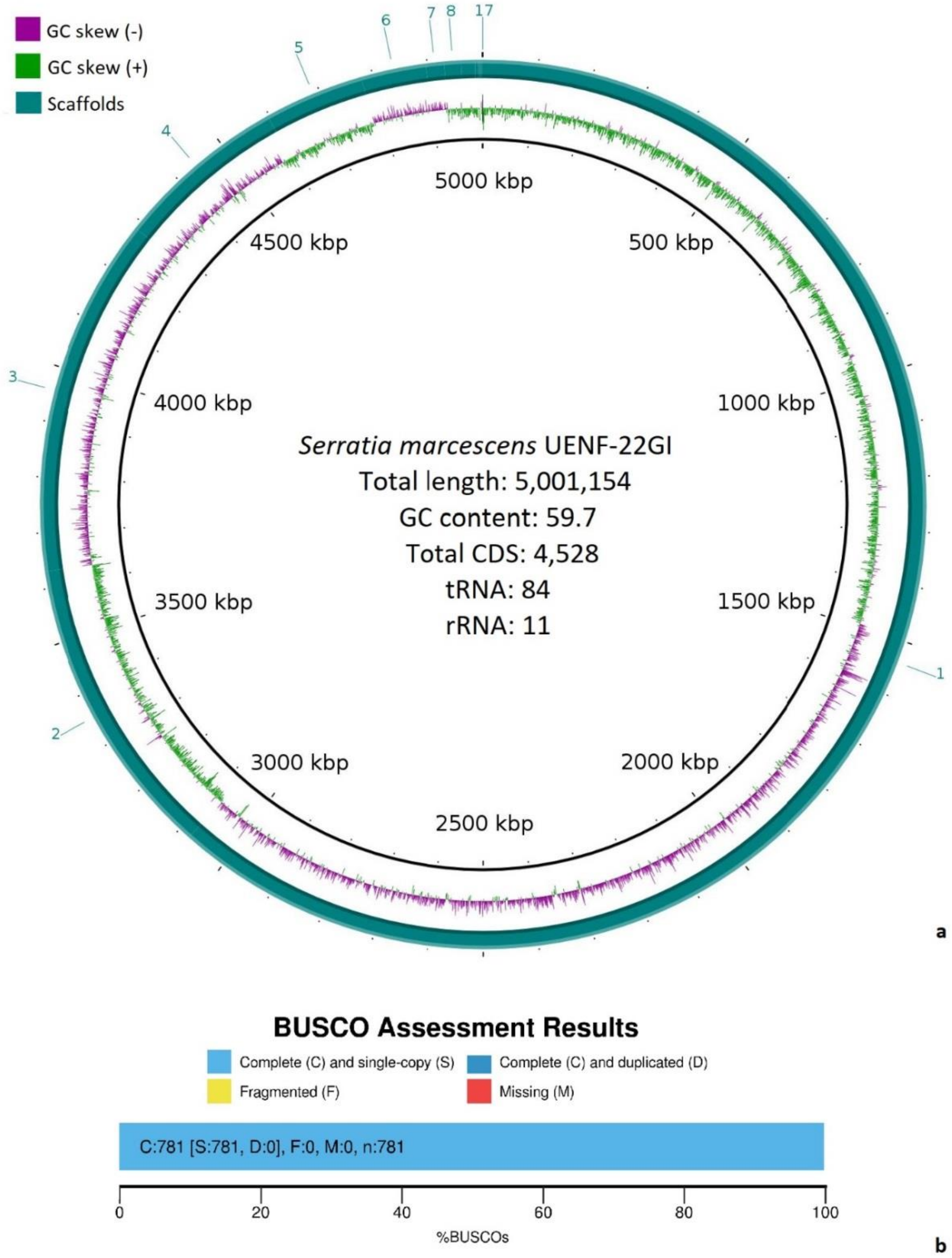


**Figure S2:** General genomic features of SMU. **(a)** Total length, number of protein-coding, tRNA and rRNA genes are represented, as well as the GC skew across the genome; **(b)** BUSCO genome completeness assessment using 781 single-copy genes from the *Enterobacteriales* reference dataset.

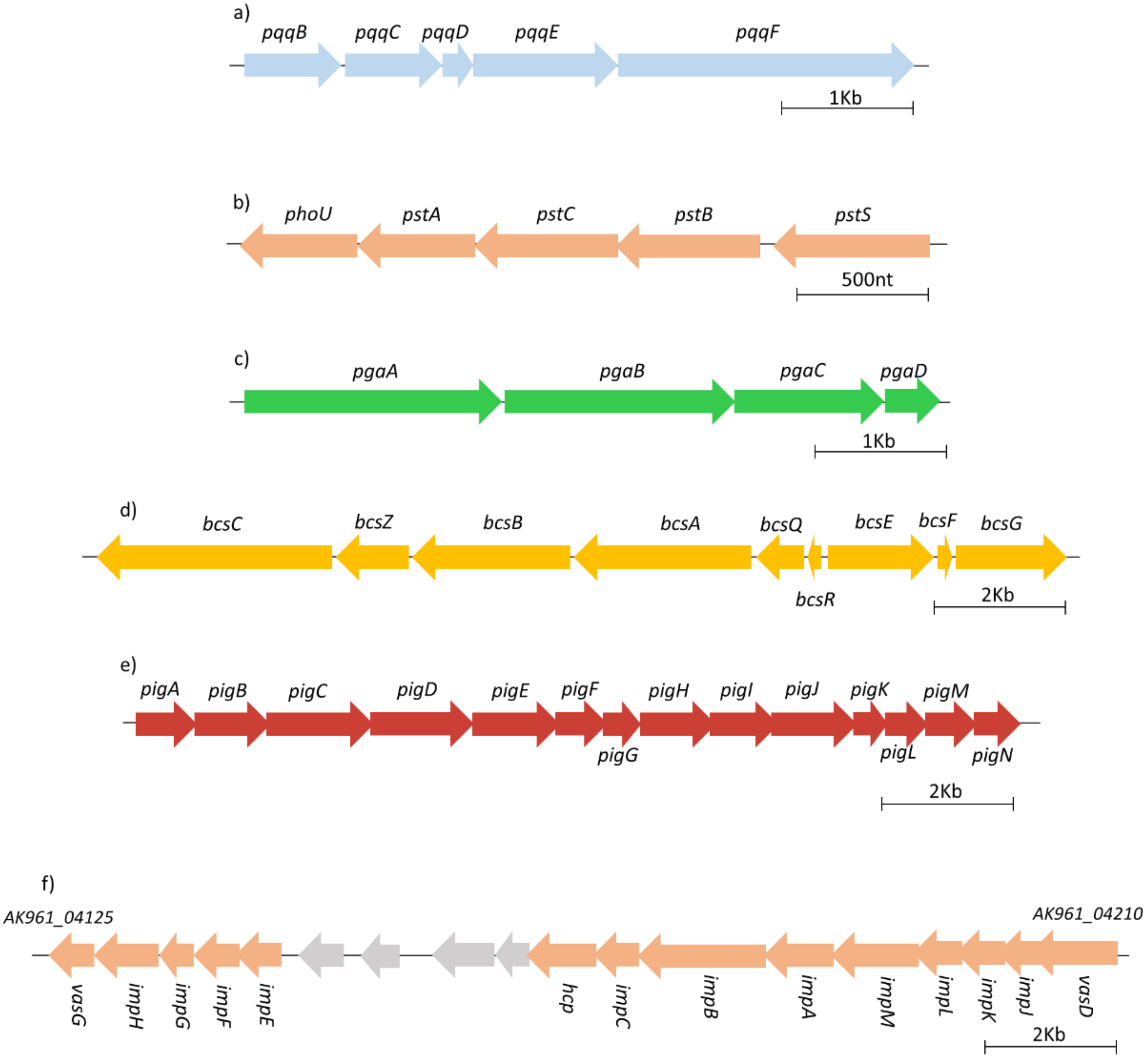


**Figure S3:** Plant growth-promoting operons found in the SMU genome. (a) biosynthesis of pqq; (b) phosphate transport system; (c) poly-beta-1,6-N-acetyl-glucosamine biosynthesis; (d) bacterial cellulose biosynthesis; (e) prodigiosin biosynthesis; (f) type VI secretion system (genes of unknown functions are in gray).

**Table S1:** Genome IDs used to build the S. marcescens pan-genome.

| Strain names | IDs |
|---|---|
| *S. marcescens* Db11 | NZ_HG326223.1 |
| *S. marcescens* WW4 | NC_020211.1 |
| *S. marcescens* FDAARGOS_65 | NZ_CP026050.1 |
| *S. marcescens* SM39 | NZ_AP013063.1 |
| *S. marcescens* CAV1492 | NZ_CP011642.1 |
| *S. marcescens* RSC-14 | NZ_CP012639.1 |
| *S. marcescens* SmUNAM836 | NZ_CP012685.1 |
| *S. marcescens* B3R3 | NZ_CP013046.2 |
| *S. marcescens* UMH1 | NZ_CP018915.1 |
| *S. marcescens* UMH2 | NZ_CP018924.1 |
| *S. marcescens* UMH3 | NZ_CP018925.1 |
| *S. marcescens* UMH5 | NZ_CP018917.1 |
| *S. marcescens* UMH6 | NZ_CP018926.1 |
| *S. marcescens* UMH7 | NZ_CP018919.1 |
| *S. marcescens* UMH8 | NZ_CP018927.1 |
| *S. marcescens* UMH9 | NZ_CP018923.1 |
| *S. marcescens* UMH10 | NZ_CP018928.1 |
| *S. marcescens* UMH11 | NZ_CP018929.1 |
| *S. marcescens* UMH12 | NZ_CP018930.1 |
| *S. marcescens* SMB2099 | NZ_HG738868.1 |
| *S. marcescens* 1274 | NZ_CP019927.1 |
| *S. marcescens* S217 | NZ_CP021984.1 |
| *S. marcescens* AS1 | NZ_CP010584.1 |
| *S. marcescens* MSU97 | GCA_001902635.1 |
| *S. marcescens* ATCC14041 | GCA_000695485.1 |
| *S. marcescens* ATCC13880 | GCA_000735445.1 |
| *S. marcescens* VGH107 | GCA_000342205.1 |
| *S. marcescens* EGD-HP20 | GCA_002265665.1 |
| *S. marcescens* CDC_813-60 | GCA_000743395.1 |
| *S. marcescens* LCT-262 | GCA_000442375.1 |
| *S. marcescens* LCT-213 | GCA_000264275.1 |
| *S. marcescens* LCT-166 | GCA_000442455.1 |
| *S. marcescens* SM39 | NZ_AP013063.1 |
| *Serratia* sp. FS14 | NZ_CP005927.1 |
| *Serratia* sp. JKS296 | GCA_900215445.1 |

**Table S2:** Unique SMU genes identified in the pan-genome analysis.

| Gene ID (NCBI PGAP) |
|---|
| AK961_02860\|hypothetical protein\|611293..613449 |
| AK961_03580\|NTPase KAP\|774541..776478 |
| AK961_03575\|hypothetical protein\|772402..774210 |
| AK961_15440\|modification methylase\|218115..219644 |
| AK961_11425\|histidine kinase\|2469256..2470674 |
| AK961_05675\|hypothetical protein\|1216374..1217663 |
| AK961_03610\|integrase\|781425..782627 |
| AK961_14170\|hypothetical protein\|3024419..3025516 |
| AK961_09800\|hypothetical protein\|2105262..2106356 |
| AK961_14195\|hypothetical protein\|3029558..3030601 |
| AK961_05655\|hypothetical protein\|1213376..1214368 |
| AK961_14205\|hypothetical protein\|3032035..3032964 |
| AK961_14190\|abortive phage resistance protein\|3028550..3029467 |
| AK961_07350\|phenazine biosynthesis protein PhzF family\|1589864..1590760 |
| AK961_10570\|N-acetylmuramoyl-L-alanine amidase\|2280871..2281728 |
| AK961_00405\|hypothetical protein\|76964..77758 |
| AK961_15630\|energy transducer TonB\|254934..255680 |
| AK961_20655\|chemotaxis protein CheY\|168079..168777 |
| AK961_07360\|hypothetical protein\|1591785..1592474 |
| AK961_10700\|hypothetical protein\|2305301..2305954 |
| AK961_14200\|resolvase\|3030972..3031595 |
| AK961_03595\|hypothetical protein\|779550..780140 |
| AK961_10695\|phage repressor protein\|2303692..2304261 |
| AK961_20680\|hypothetical protein\|176271..176828 |
| AK961_03585\|hypothetical protein\|776776..777321 |
| AK961_05670\|hypothetical protein\|1215167..1215676 |
| AK961_14625\|pilus assembly protein\|43995..44489 |
| AK961_02105\|hypothetical protein\|436034..436510 |
| AK961_05660\|hypothetical protein\|1214474..1214941 |
| AK961_05650\|hypothetical protein\|1212788..1213231 |
| AK961_15465\|hypothetical protein\|224687..225112 |
| AK961_22105\|hypothetical protein\|177855..178256 |
| AK961_03605\|hypothetical protein\|781042..781422 |
| AK961_09475\|alpha/beta hydrolase\|2042223..2042588 |
| AK961_10685\|hypothetical protein\|2301158..2301517 |
| AK961_02320\|hypothetical protein\|485474..485833 |
| AK961_10370\|hypothetical protein\|2228241..2228588 |
| AK961_03490\|hypothetical protein\|743718..744053 |

AK961_14585|hypothetical protein|36315..36644
AK961_00400|addiction module toxin RelE|76300..76629
AK961_22995|hypothetical protein|3653..3967
AK961_11400|hypothetical protein|2465145..2465450
AK961_09560|hypothetical protein|2057145..2057450
AK961_00135|hypothetical protein|32913..33179
AK961_20740|hypothetical protein|189167..189430
AK961_09805|hypothetical protein|2106476..2106736
AK961_05870|hypothetical protein|1257201..1257452
AK961_20735|hypothetical protein|188925..189170
AK961_03590|hypothetical protein|779307..779549
AK961_14210|hypothetical protein|3033883..3034098
AK961_03600|hypothetical protein|780494..780709
AK961_05665|hypothetical protein|1214955..1215170
AK961_01290|hypothetical protein|273026..273232
AK961_08365|hypothetical protein|1817511..1817699
AK961_03570|hypothetical protein|772181..772366
AK961_11035|hypothetical protein|2383022..2383204
AK961_17755|hypothetical protein|180397..180576
